# Dendritic Cell – Fibroblast Crosstalk via TLR9 and AHR Signaling Drives Lung Fibrogenesis

**DOI:** 10.1101/2024.03.15.584457

**Authors:** Hannah Carter, Rita Medina Costa, Taylor S. Adams, Talon Gilchrist, Claire E. Emch, Monica Bame, Justin M. Oldham, Angela L. Linderholm, Imre Noth, Naftali Kaminski, Bethany B. Moore, Stephen J. Gurczynski

**Author notes:** Corresponding author: Stephen J. Gurczynski, PhD 5608C Medical Science II 1137 Catherine St Ann Arbor, MI 48109-2200. The Authors declare no conflicts of interest. **Funding sources.** This work was funded by NIH R35 HL144481 awarded to BBM, T32GM007863 awarded to HC, and a Boehringer Ingelheim discovery award in ILD. This work was made possible by the Pulmonary Fibrosis Foundation, by an independent grant from Boehringer Ingelheim Pharmaceuticals, Inc. who provided the financial support. The authors meet criteria for authorship as recommended by the International Committee of Medical Journal Editors (ICMJE) and were fully responsible for all aspects of the trial and publication development. **Author contributions.** HC performed experiments, analyzed data, wrote, and edited the manuscript. TSA analyzed and interpreted data. RMC, TG, CE, and MB performed experiments and analyzed data. JMO, ALL, and IN provided clinical data. BBM analyzed and interpreted data and edited the manuscript. SJG wrote and edited the manuscript as well as analyzing and interpreting data.

## Abstract

Idiopathic pulmonary fibrosis (IPF) is characterized by progressive scarring and loss of lung function. With limited treatment options, patients succumb to the disease within 2-5 years. The molecular pathogenesis of IPF regarding the immunologic changes that occur is poorly understood. We characterize a role for non-canonical aryl-hydrocarbon receptor signaling (ncAHR) in dendritic cells (DCs) that leads to production of IL-6 and IL-17, promoting fibrosis. TLR9 signaling in myofibroblasts is shown to regulate production of TDO2 which converts tryptophan into the endogenous AHR ligand kynurenine. Mice with augmented ncAHR signaling were created by crossing floxed AHR exon-2 deletion mice (AHR_Δex2_) with mice harboring a CD11c-Cre. Bleomycin was used to study fibrotic pathogenesis. Isolated CD11c+ cells and primary fibroblasts were treated ex-vivo with relevant TLR agonists and AHR modulating compounds to study how AHR signaling influenced inflammatory cytokine production. Human datasets were also interrogated. Inhibition of all AHR signaling rescued fibrosis, however, AHR_Δex2_ mice treated with bleomycin developed more fibrosis and DCs from these mice were hyperinflammatory and profibrotic upon adoptive transfer. Treatment of fibrotic fibroblasts with TLR9 agonist increased expression of TDO2. Study of human samples corroborate the relevance of these findings in IPF patients. We also, for the first time, identify that AHR exon-2 floxed mice retain capacity for ncAHR signaling.

## Introduction

Idiopathic Pulmonary Fibrosis (IPF) is a progressive, fibrotic disease with estimated survival of 2-5 years after diagnosis. Risk factors include male sex, cigarette smoking, repeated viral infection, with median age at diagnosis of 66 years.(1) Treatments slow progressive fibrosis, but only lung transplant is curative.(2) Fibroblast activation and excess production of collagens are critical factors that underlie disease progression, however, fibrogenesis is a complex and poorly understood process requiring cooperation / crosstalk between multiple cell subsets. Myriad inflammatory cells accumulate in fibrotic lungs including dendritic cells (DCs), macrophages, and lymphocytes, however, the ultimate role of the immune system in initiation and progression of fibrosis is not understood. Immune responses may promote progressive scarring but treating patients with steroids is counterintuitively harmful highlighting the need for more research into the interactions between immune cell populations and fibrogenesis.(3) We have previously defined an important role for tissue resident DCs, and more specifically DCs expressing the aryl-hydrocarbon receptor (AHR) in viral-mediated pulmonary fibrosis following bone marrow transplantation.(4, 5) Tissue resident DCs have been shown to accumulate in the lungs of both IPF patients and bleomycin treated mice, however their exact role in the initiation or progression of pulmonary fibrosis is unclear. (6–8)

The canonical Aryl-Hydrocarbon Receptor (AHR) pathway is an intracellular signaling cascade originally described as the major sensor of a poly-cyclic aromatic hydrocarbon (PAH) toxicants, e.g., dioxin.(9) More recently, endogenous AHR ligands such as the kynurenine family of tryptophan metabolites have been described.(10, 11) Kynurenine can be made by indoleamine-2,3-dioxygenase (IDO) or tryptophan 2,3-dioxygenase (TDO), and kynurenine / tryptophan ratios are increased in patients with fibrotic lung disease.(12) Increased kynurenine / tryptophan ratios in the blood are also prognostic indicators for many inflammatory lung diseases, including cancers and SARS-CoV-2.(13–18)

AHR is constitutively expressed in an inactive form sequestered in the cytoplasm bound to various chaperone proteins.(19) Binding of ligand exposes a nuclear localization signal which facilitates translocation of AHR to the nucleus wherein the chaperone coat is recycled and AHR dimerizes with its major canonical partner ARNT.(20, 21) This AHR / ARNT dimer binds to xenobiotic response elements in the genome and activates transcription of a variety of gene products including the CYP family of cytochrome p450 oxidases.(22)

AHR is present and involved in inflammation in several lung cell types including immune cells such as DCs, macrophages, and lymphocytes as well as structural cells such as epithelial cells and fibroblasts. Recently AHR has been studied for its interactions with other binding partners outside of ARNT, i.e., NFkB subunits RelA / RelB and KLF6, in what is termed “non-canonical AHR signaling” (ncAHR) which is especially important in regulating expression of pro-inflammatory cytokines in immune cells like dendritic cells (DCs) and macrophages.(4, 23–25) Production of pro-inflammatory cytokines by DCs is augmented in the presence of AHR ligands suggesting a role for AHR in concert with NFkB when triggered by pro-inflammatory signals.(26, 27) AHR and TLR signaling, especially TLR9, have also been studied separately for their ability to promote fibrogenesis by directly augmenting fibroblast differentiation into pro-fibrotic myofibroblasts.(28, 29) Pro-inflammatory TLR9 / AHR responsiveness has been linked to worsened IPF progression(30), however, the functional binding partners of AHR in inflammatory vs normal conditions, the triggers which divert AHR from its canonical pathway, and the exact role of AHR in complex pathologies such as fibrosis are not well understood.

Here, we present data that ncAHR signaling is an important modulator of DC immune responses during the progression of pulmonary fibrosis. Using the bleomycin mouse model of pulmonary fibrosis, we demonstrate that AHR+ DCs accumulate in the lungs and drive production of IL-6 in an AHR dependent fashion. AHR plays a role in differentiation of several immune cells, including Treg, Th-17 and Th-22 cells(31–33), which makes whole-body knock outs non-ideal for studying immune responses. Using a DC-specific deletion of the DNA binding region of AHR (CD11c-Cre / AHR_Δex2_), we further demonstrate deletion of canonical AHR function in DCs leads to significant upregulation of ncAHR signaling, pro-inflammatory cytokines and worsened fibrosis in the bleomycin model. This work also demonstrated that adoptive transfer of ncAHR-activated DCs can promote fibrogenesis in a CD4 T cell-independent manner, highlighting an unappreciated role for DCs in fibrogenesis. We also provide evidence that myofibroblasts in the lungs of both IPF patients and fibrotic mice express the enzymes that convert tryptophan into endogenous AHR ligands highlighting an important role for TDO2 specifically in the fibrogenic circuit that is established between structural cells and immune cells in the fibrotic lung.

## Materials and Methods

### Mouse line, cell lines, and reagents

All experiments were conducted under an IACUC approved animal protocol. WT C57Bl/6J mice were ordered from Jackson labs and used between 6-8 weeks of age. AHR_Δex2_ mice were generated by crossing the AHR^fx^ line (Ahrtm3.1Bra/J, strain 6203, Jackson labs) with a CD11c-Cre expressing line (B6.Cg-Tg(Itgax-cre)1-1Reiz/J, strain 8068, Jackson labs). Resulting pups were verified to contain both Cre and the AHR floxed alleles via PCR using Jackson labs recommended primers. CD11c+ cells were further verified to contain the AHR exon2 excised allele via PCR using Jackson labs recommended primers. AHR_Δex2_ mice were bred at the University of Michigan mouse breeding facility and used between 6-8 weeks of age. For some experiments AHR^fx^ mice which did not express CD11c-Cre were used as littermate controls. Bleomycin was purchased as Blenoxane from the pharmacy at the University of Michigan hospital, 1U/mL stocks were made using sterile saline and mice were administered 0.75U/kg via the oral-pharyngeal route. The AHR and TDO2 inhibitors CH223191 and 680C91 (Tocris) were used at 5 mg/kg and 15mg/kg respectively for *in vivo* experiments, CH223191 was used between 10-30 μM for *in vitro* treatment of cell culture. The TiPARP (PARP7) inhibitor RBN2397 (Selleckchem) was used at a concentration of 10 μM. The TLR9 agonist ODN2395 (Invivogen) was used between 0.1 – 1.0 μM.

### Generation of inducible iCD103+ DCs

Inducible CD103 (iCD103) DCs were generated from mouse bone marrow as previously described.(4, 34) In short, bone marrow cells were aseptically isolated from the femurs of indicated mouse lines and incubated for 16-days in RPMI media containing 10% heat inactivated FBS, 1% Pen/Strep cocktail, 50µM β-mercapto-ethanol, 3 ng/mL GMCSF (R&D systems) and 200 ng/mL Flt3-L (R&D systems) before any additional treatment. iCD103 cells were characterized to be CD11c+, MHCII+, and CD103+ by flow cytometry. For adoptive transfer experiments, Groups of WT-B6 mice were first administered BLM before receiving 1x10^6^ iCD103+ cells from AHR_Δex2_ mice via tail vein injection. CD4 T-cell depletion was carried out via i.p. injection of 100 μg anti-CD4 depleting antibody (clone GK1.5 Bioxcell) at 0- and 7-days post BLM instillation.(35)

### Fibroblast isolation

Groups of mice were administered BLM or saline as control. 21-days post BLM, lungs were harvested and minced with scissors. Minced lungs were incubated in complete DMEM (DMEM + 10% FBS and 1% pen/strep cocktail) for 7 days to let fibroblasts adhere.

Media was changed at 7 days and every 2 days afterwards. Cells were trypsonized and plated on day 14-post isolation for subsequent assays.

### Single Cell RNA-seq analysis

Processed gene expression data from Habermann et al.(36) was downloaded from GSE135893. Raw sequencing data from Adams et al.(37) was downloaded from GSE136831 for reprocessing. Read 2 poly(A) and template-switch oligo contaminants were trimmed using cutadapt (version 3.1). Reads were aligned to genome GRCh38 with STAR (version 2.7.8a) using Gencode annotation release 38.

Control and IPF cells previously classified as stromal were filtered from each dataset and combined in R (version 4.0.3) using the library Seurat (version 4.3.0.1). For normalization, raw expression counts were scaled to 10,000 transcripts per cell then natural log transformed with a pseudocount of 1. Cells were split by dataset and integrated using Seurat’s reciprocal principal component analysis (RPCA) implementation, followed by dimension reduction, graph embedding, high granularity clustering and cell type re-annotation. These steps were performed iteratively until all multiplets were identified and removed.

Of the filtered data presented: Habermann et al. consists of 2,801 cells from 10 controls and 12 IPF lungs; Adams et al. consists of 6,850 cells from 26 control and 32 IPF lungs. Integrated gene expression values were only used for PCA-based graph embedding; cluster interrogation and subsequent visualizations were performed with normalized, empirical values. For the heatmap, normalized gene expression values of each cell type were averaged for each subject with 3 or more respective cells. To avoid batch effect between datasets when visualizing gene expression, normalized expression values were further min-max normalized across cells within each dataset.

### Generation of single cell suspensions and flow cytometry

Single cells suspensions were generated via collagenase digestion of lung tissue as previously described.(38) In short, mouse lungs were first perfused via injection of 3-5mL of saline through the right ventricle of the heart. Lungs were resected and minced with scissors before incubation in a collagenase buffer containing 1 mg/mL Collagenase-A (Roche), 2000U DNaseI (Sigma), in complete DMEM media supplemented with 10% heat inactivated FBS and 1% Pen/Strep cocktail for 45 minutes at 37°C in a shaking incubator. Cell suspensions were disrupted by drawing through a 10 mL syringe before filtration through a 100 µm pore sized cell strainer. Resulting single cell suspensions were assayed for cell viability via trypan blue exclusion, staining for flow cytometry was subsequently carried out via incubation with appropriately diluted fluorophore conjugated antibodies. Antibodies used for flow cytometry were as follows: Myeloid flow cytometry staining panel, BV650-CD11b (clone M1/70), BV510-CD45 (clone 30-F11), BV421-I-Ab (MHCII clone AF6-120.1), APC-Cy7-SiglecF (clone E50-2440), APC-Ly6G (clone 1A8) purchased form BD Horizon. BV605-CD64 (clone X54-5/7.1), PerCP-Cy5.5-CD24 (clone M1/69), PE-Cy7-Ly6C (clone HK1.4), PE-CD193 (CCR3 clone J073E5) purchased from Biolegend. PE-eFluor610-CD11c (clone N418), PE-AHR (clone 4MEJJ) purchased from eBioscience. Lymphocyte flow cytometry staining panel. PE-Cy7-IFNγ (clone XMG1.2), FITC-CD3 (clone 17A2) purchased from BD bioscience. APC-Cy7-IL17A (clone TC11-18H10.1), BV570-CD8a (clone 53-6.7), AF700-CD90.2 (Thy1.2 clone 30-H12), BV510-CD4 (clone GK1.5) purchased from Biolegend. PE-eFluor 610-FoxP3 (clone FJK-16s) purchased from eBioscience. Note: Some markers were stained for but were not included in this study. Flow analysis was carried out in FlowJo v. 10.5.

### Quantification of collagen via hydroxyproline assay

Collagen was quantified as described in(39). In short, lungs from BLM or WT mice were resected at the indicated timepoints and homogenized in a buffer of PBS and complete protease inhibitor (Roche). Lung homogenates were diluted 1:1 with 12N hydrochloric acid and incubated overnight at 120°C in well-sealed glass tubes. The following day, aliquots of acid-hydrolyzed lung homogenate were incubated with a chloramine-T solution and then reacted with 4-(Dimethylamino)benzaldehyde.

Hydroxyproline was detected via absorbance at 560 nM on a synergy H1 spectrophotometer (Biotek).

### Isolation of RNA for qRT-PCR

RNA was purified from lung tissue or cell pellets via incubation with Trizol reagent following manufacturer directions. RNA quantity and purity was assessed on a nanodrop spectrophotometer (Thermo). Transcript expression was analyzed via reverse transcriptase quantitative real-time PCR using Lunaprobe reagent (New England Biolabs) on a Quantstudio3 thermocycler (ABI). Expression analysis was conducted using a ΔΔct calculation. Transcripts were normalized to expression level of a housekeeping gene, RPL38, using a pre- validated primer probe set (Integrated DNA technologies, Hs.PT.58.40595235.gs). Other primer and probe sequences are detailed in Table 1.

### Analysis of plasma concentrations of IL-6 and IL-17 isoforms

Circulating plasma concentration for IL6 and IL17 (a, c, d and f isoforms) was determined used the Olink (Uppsala, Sweden) Explore platform, which uses proximity extension assays to generate semi-quantitative protein data, which are Log2 transformed for modeling. The association between plasma biomarker concentration and three-year transplant-free survival was assessed in a combined cohort of patients with IPF (n=366) from the University of Virginia, University of Chicago and University of California at Davis using univariable Cox proportional hazards regression [IRB approval and subject characteristics were previously reported(40)]. Survival was plotted for each quartile of IL6 concentration using the Kaplan-Meier estimator. The proportional hazards assumption was checked and satisfied for each biomarker tested.

### Statistical analysis

Statistical analysis was conducted in Graphpad Prism v.9.1.0. Statistical significance was determined using students T-test when two groups were compared and ANOVA using Holm-Sidak’s post-test when three or more groups were compared. Relevant statistical information, i.e., P-values and n-values, are given in individual figure legends.

## Results

### CD103+ DCs accumulate in the fibrotic lung, promote fibrogenesis and show evidence of pro-inflammatory AHR signaling

We previously published that AHR+ CD103+ DCs were recruited to lungs, and drove lung injury and fibrosis following bone marrow transplantation and herpesvirus infection(4); however, a pathogenic role for AHR and DCs in non-infectious forms of pulmonary fibrosis is not known. To address this, we characterized lung DCs using a bleomycin model of pulmonary fibrosis. Lung leukocytes were analyzed via flow cytometry at 7- and 14-days post-treatment and a significant increase in CD103+ DCs (CD45+, CD11c+, MHCII +, CD103+, CD11b-) was noted at both time points (Fig. 1A). Differentiation of CD103+ DCs requires the transcription factor BATF3 and CD103+ DCs are completely absent in BATF3^-/-^ mice.(41) To clarify the role of lung CD103+ DCs in mediating fibrogenesis we treated BATF3^-/-^ mice with bleomycin for 14-21 days and verified the absence of CD103+ DCs in the lung by flow cytometry (Fig. 1B). BATF3^-/-^ mice exhibited durable protection from bleomycin induced pulmonary fibrosis exhibiting less collagen production at 14 days as assessed by hydroxyproline assay and at 21-days as assessed by qRT-PCR for collagen I transcript (Fig. 1C and 1D).

**Figure 1.**
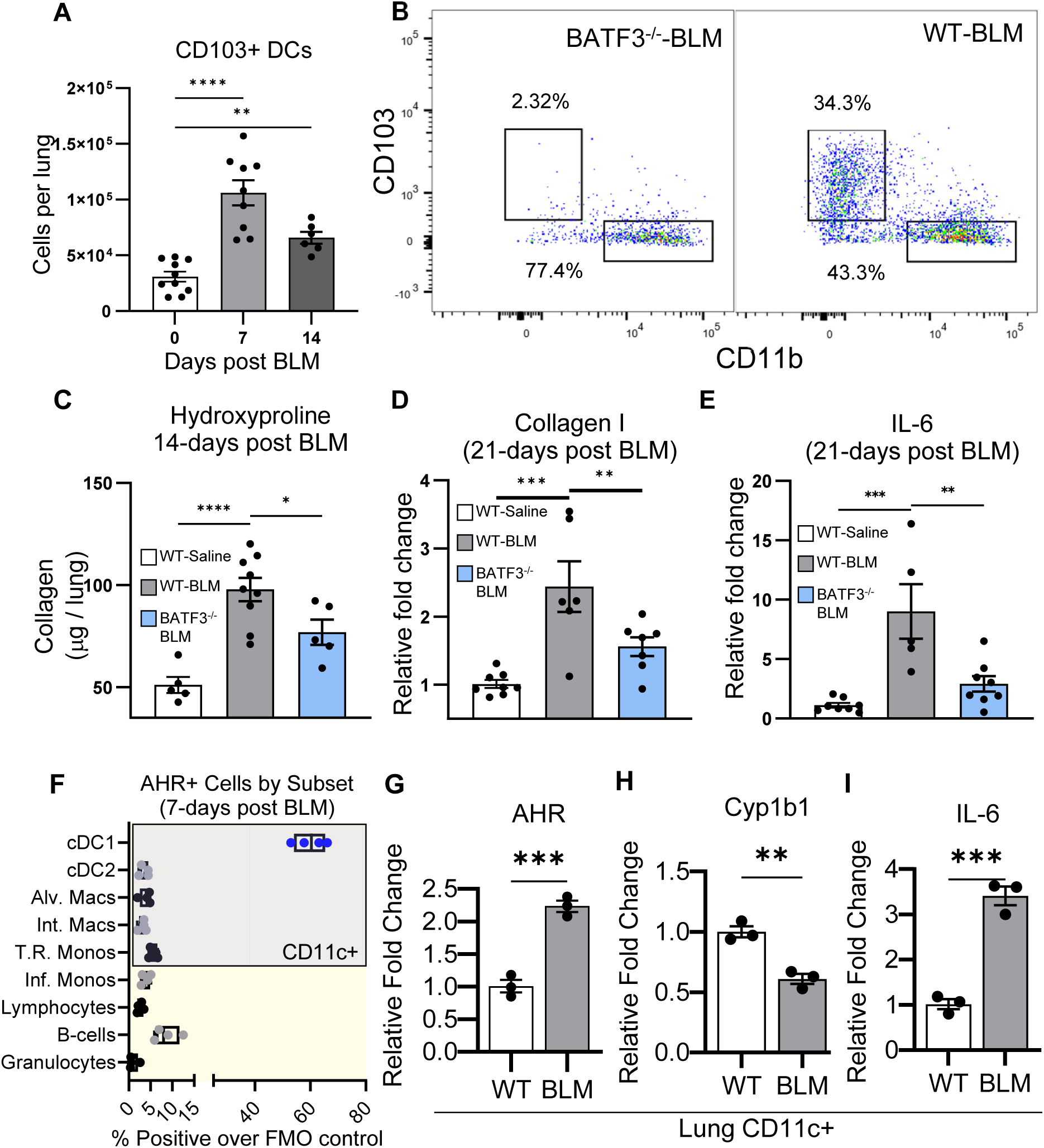
CD103+ DCs accumulate in the fibrotic lung and show evidence of pro-inflammatory AHR signaling. **A and B)** Mice (n = 6 – 10 mice per group (panel A), 5 per group (panel B) were treated with 0 75 U/kg bleomycin for 0, 7 or 14 days (panel A) or 14 days (panel B). Lungs were harvested and single cell suspensions were analyzed by flow cytometry for the presence of CD103+ DCs (CD45+, CD11c+, MHCII+, CD24+, CD64-, CD103+). **(C, D, and E)** groups of mice (n = 5-8 mice per group (panel C) or n = 6-8 per group (panel D and E)) were treated with 0.75 U/kg BLM for 14 – 21 days. Lungs were harvested and collagen was quantified by hydroxyproline assay or qRT-PCR. **F)** Mice (n = 4 per group) were treated with 0.75 U/kg BLM for 7 days after which lungs were harvested and single cell suspensions were analyzed via flow cytometry for AHR expression in the indicated cell subsets. **G through I)** Mice (n = 5 per group) were treated with 0.75 U/kg bleomycin or saline control (WT). Single cell suspensions were prepared from collagenase digested lung tissue at 7-days post bleomycin and CD11c+ cells were purified via magnetic bead isolation. Expression of the indicated transcripts was analyzed via qRT-PCR. All experiments shown are representative of at least three independent experiments, statistical significance was determined via ANOVA in panel A and C-E, or student’s t-test in panels G-I (* = p < 0.05, ** = p < 0.01, *** = p < 0.001, **** = p < 0.0001).

Interestingly, BATF3^-/-^ mice also displayed less IL-6 transcript indicating that CD103+ DCs are important regulators of pro-inflammatory / fibrotic cytokine production following bleomycin challenge (Fig. 1E). We next analyzed expression of AHR in various lung immune cells via flow cytometry at 7-days post bleomycin treatment (markers for cell subsets are detailed in Sup. Table 1). The highest percentage of AHR+ cells was found in CD103+ DCs (cDC1s ∼60%) while most other immune cells had AHR+ rates between 3-5% (Fig. 1F). To further verify that expression of AHR was augmented in lung DCs we isolated CD11c+ cells via magnetic bead isolation from the lungs of bleomycin treated or control mice. CD11c+ cells from the lungs of bleomycin treated mice had increased expression of AHR in comparison to control mice but interestingly had decreased expression of the canonical AHR gene product Cyp1b1 (Fig. 1G and 1H). In agreement with our previous data using BATF3^-/-^ mice, CD11c+ cells isolated from bleomycin treated mice had increased expression of IL-6 further indicating that lung DCs are an important source of pro-inflammatory cytokines (Fig. 1I).

### Production of AHR ligands is augmented during BLM-induced fibrosis and complete AHR inhibition ameliorates disease

Several endogenous ligands for AHR have been described, most notably, kynurenine which is a metabolite of tryptophan processed by the heme-dependent oxygenases IDO1/2 and TDO2.(42) Whole lung homogenates were prepared from saline-treated or BLM-instilled mice. Concentrations of lung-associated kynurenine were assayed via ELISA. A significant increase in lung-associated kynurenine was observed at both 7- and 14-days post-BLM (Fig. 2A). These data suggest activation of AHR signaling in the fibrotic lung may promote development of BLM-induced fibrosis. To test this hypothesis, we administered the complete AHR inhibitor, CH223191, following BLM instillation. Complete inhibition of AHR signaling ameliorated development of bleomycin-induced pulmonary fibrosis. Mice treated with CH223191 exhibited less collagen deposition than non-treated mice (Fig. 2B) and had decreased evidence of fibrotic pathology on trichrome stained lung sections (Fig. 2C).

**Figure 2.**
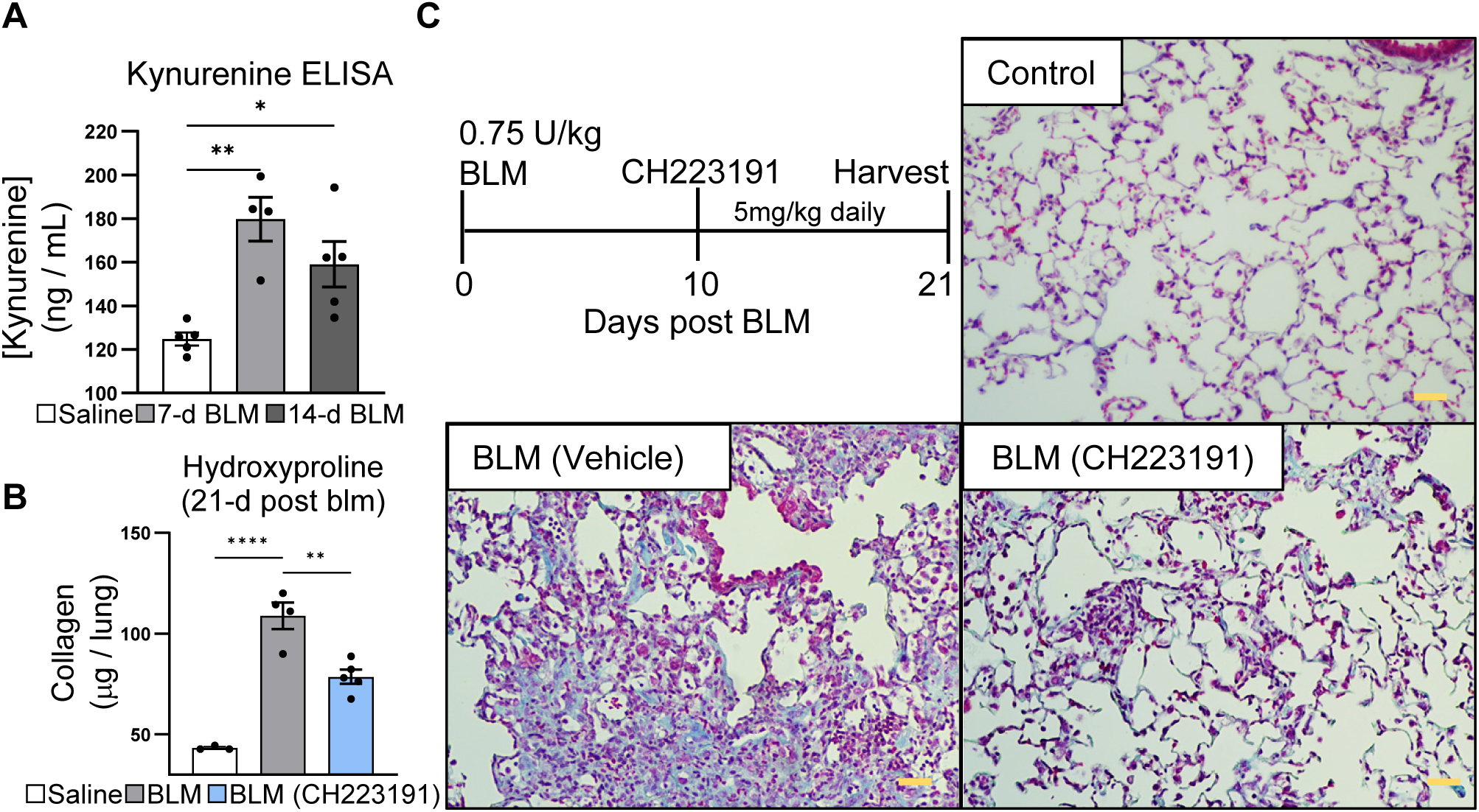
Production of AHR ligands is augmented during BLM induced fibrosis and complete AHR inhibition ameliorates disease. **A)** Mice (n = 4-5 mice per group) were treated with 0.75 U/kg BLM. Lungs were harvested at the indicated times and whole lung homogenates were assayed for KYN concentration by ELISA. **B)** Mice, (n = 3 – 5 per group) were treated with 0.75 U/kg bleomycin. Beginning at day 10 mice were treated daily with 5 mg/kg CH223191 dissolved in a vehicle of 2% Tween-20, 0.5% Carboxy methylcellulose, in PBS or vehicle alone by oral gavage until day 18. Lungs were harvested at Day 21 and lung collagen was quantified by hydroxyproline assay. **C)** Representative histology of lungs from B). Sections were stained with trichrome which highlights collagen fibers as blue staining (scale bar = 50 μM). All experiments shown are representative of at least three independent experiments, statistical significance was determined via ANOVA (* = p < 0.05, ** = p < 0.01, **** = p < 0.0001).

### Deletion of the AHR DNA binding domain reduces canonical AHR signaling in DCs but AHR expression is retained and augments inflammatory non-canonical AHR signaling

To further address AHR signaling in lung DCs we bred mice that harbored floxed copies of AHR exon-2 (43) crossed to a strain that harbored Cre recombinase under control of the CD11c promoter (CD11c-AHR_Δex2_ mice referred to as AHR _Δex2_, Fig. 3A). Previous reports using these AHR exon-2 floxed mice described them as AHR-null mutants (43); however, we identified an alternative start codon (Supplemental Fig. 1A red text) that is introduced as part of the fusion of AHR exon-1 and exon-3 resulting in a 14 amino acid N-terminal leader sequence (Supplemental Fig. 1A green text) fused in frame to AHR exons 3 – 10 (Supplemental Fig. 1A blue text). DCs generated from these mice are indeed null for production of canonical AHR response genes following stimulation with kynurenine (Fig. 3B), however, primers specific for AHR exon3-4 showed no difference in the amounts of AHR transcript while primers specific for the junction of exon1-2 showed a significant reduction in AHR_Δex2_ mice (Fig. 3C). Furthermore, AHR was still detectable in CD103+ DCs produced from AHR_Δex2_ mice as assessed by intracellular flow cytometry (Fig. 3D and 3E).

**Figure 3.**
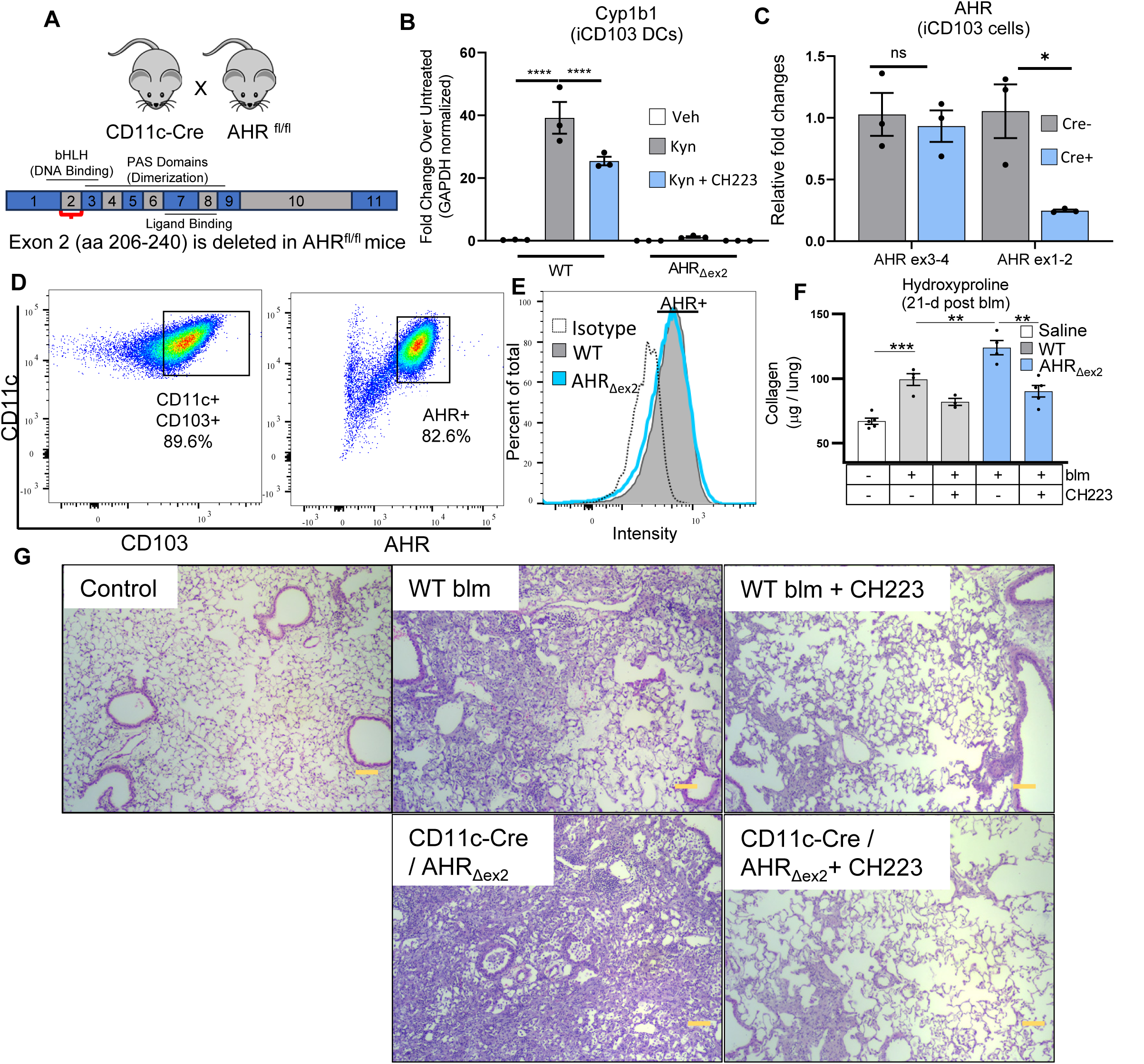
Deletion of AHR exon-2 reduces canonical AHR signaling in DCs but AHR expression is retained and augments inflammatory non-canonical AHR signaling. **A)** Cartoon schematic of murine AHR depicting the deletion of AHR exon-2 containing the DNA binding bHLH domain. Numbered boxes correlate to individual exons. **(B and C)** iCD103+ DCs were generated from WT B6 mice or AHR_Δex2_ mice and treated with 200 μM KYN or 10 μM CH223191 (CH223). 18 hours after treatment, RNA was harvested and expression of the canonical AHR gene Cyp1b1 (A) or AHR (B) was analyzed via qRT-PCR. **D)** iCD103+ DCs were analyzed via flow cytometry for expression of CD11c and CD103 as well as intracellular expression of AHR. **E)** Representative histogram showing AHR expression in WT or AHR_Δex2_ iCD103 DCs analyzed via flow cytometry. **F and G)** Mice (n = 5 per group) were treated with 0.75 U/kg bleomycin. At 10-d post bleomycin groups of mice were administered CH223191 (5 mg/kg) or vehicle alone daily until 18-d post bleomycin. Lungs were harvested at 21-d post bleomycin collagen content was quantified via hydroxyproline assay (F) and lung fibrosis was examined via histopathologic examination of fixed lung sections (A: Scale bar = 100 μM panel G). All experiments shown are representative of at least three independent experiments, statistical significance was determined via ANOVA in panel A or student’s t-test in panel B (* = p < 0.05, ** = p < 0.01, **** = p < 0.0001).

We next generated cDNA from BMDCs isolated from either heterozygous B6 / AHR _Δex2_ or homozygous AHR_Δex2_ mice. PCR of the AHR ORF produced two bands in the heterozygous mice of approximately 2400 and 2200 bp (Supplemental Fig. 1B) corresponding to the predicted sizes of the full length AHR or AHR_Δex2_ ORF, respectively, and a single band of approximately 2200bp in the AHR_Δex2_ mice. We cloned these products into pl-Cherry-NEO vector which co-expressed each construct with mCherry and verified sequences of both full length and AHR_Δex2_ PCR products via sanger sequencing (data not shown) . To verify the specificity of the AHR flow cytometry antibody, We conducted flow cytometry on 3T3 cells transfected with either AHR full length or AHR_Δex2_ constructs and noted ∼30% of cells expressed mCherry after 48 hours (Supplemental Fig. 1C). We also examined expression of AHR in these transfected cells and noted a ∼2-fold increase in AHR MFI in both full length and AHR_Δex2_ transfections indicating both constructs produced an in frame AHR protein. Thus, we now report that although canonical AHR signaling is ablated following excision of exon-2, a truncated AHR protein is produced that likely mediates effects through the ncAHR pathway.

To assess impact of DC AHR signaling on development of fibrosis we administered BLM to groups of WT B6 or AHR_Δex2_ mice. AHR_Δex2_ mice exhibited increased fibrosis and ∼1.5-fold increased collagen deposition 21-days post BLM versus WT mice (Fig. 3F and 3G). We were surprised by this result as complete AHR inhibition ameliorated fibrosis (Fig. 1C). We next administered CH223191 to groups of BLM-treated WT or AHR_Δex2_ mice. Interestingly, complete inhibition of AHR was still ameliorative (Fig. 3F) indicating that, while deletion of AHR exon-2 effectively ablated canonical AHR signaling in CD11c+ cells (Fig 3B), ncAHR signaling was likely still functional and could mediate pro-fibrotic effects.

### Loss of canonical AHR signaling in CD11c+ cells exacerbates fibrosis via a ncAHR-dependent mechanism

To further test the role AHR in CD103+ DCs we generated pure populations of induced CD103+ DCs (iCD103) from bone marrow stem cells.(34) We previously published iCD103 DCs express many cDC1 markers and express AHR.(4) We addressed the role of CD103+ DCs in mediating fibrosis by adoptively transferring iCD103+ DCs from AHR_Δex2_ mice into groups of BLM treated WT mice. Lungs were harvested at 21-days post BLM and collagen content was assessed via hydroxyproline assay. B6 BLM treated mice that had received an adoptive transfer of AHR_Δex2_ iCD103+ cells developed increased fibrosis exhibiting significantly increased lung collagen over mice receiving a sham transfer (Fig. 4A). We also administered neutralizing αCD4 antibody to BLM-treated mice with AHR_Δex2_ iCD103+ adoptive transfer, however, no change in fibrosis was observed indicating the pro-fibrotic effects of AHR_Δex2_ CD11c+ cells are independent of CD4+ T-cells(Fig. 4A).

**Figure 4.**
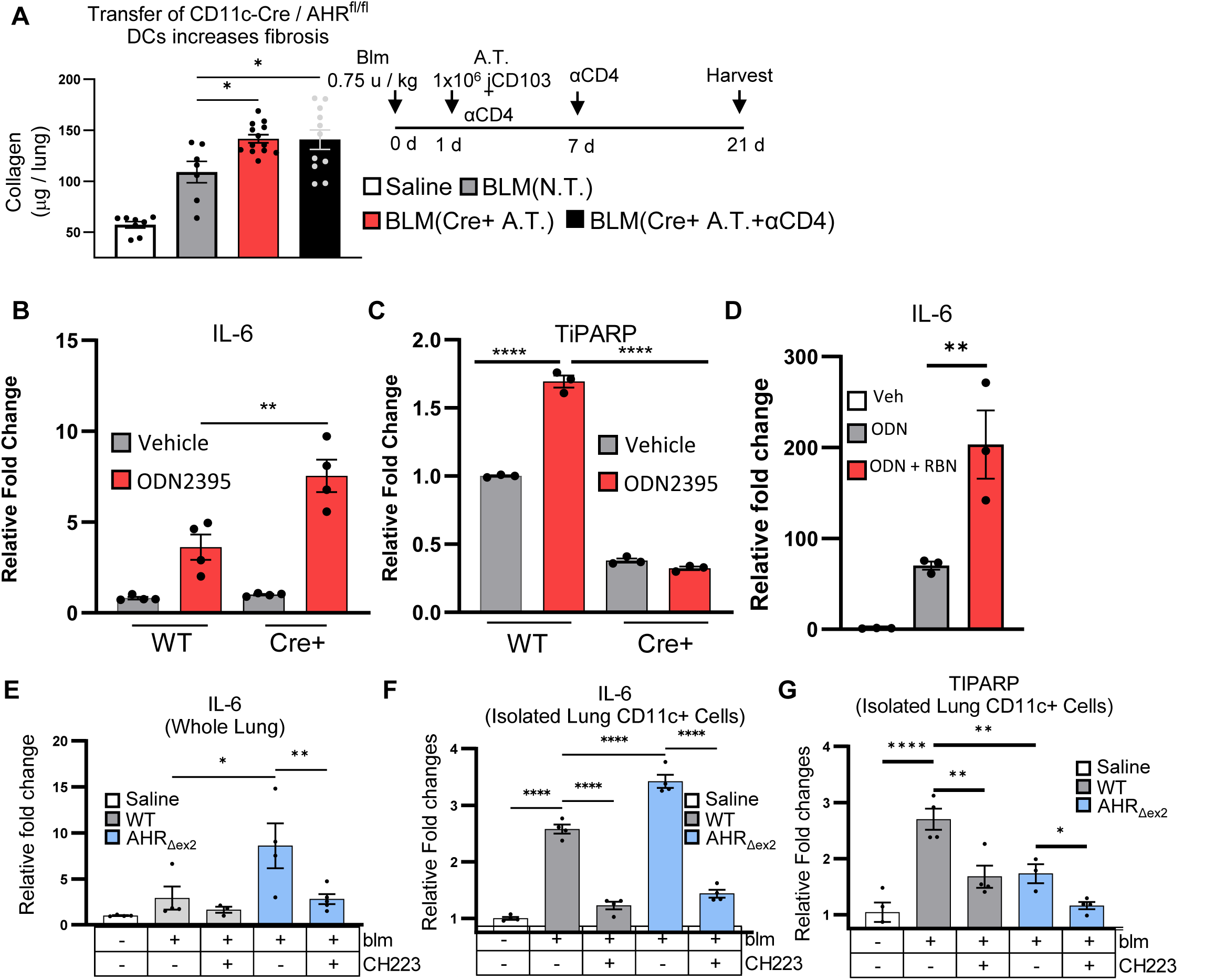
Loss of canonical AHR signaling in CD11c+ cells exacerbates fibrosis via a ncAHR dependent mechanism. **A)** Mice (n = 7 mice no transfer group and 11 mice Cre+ transfer groups) were treated with 0.75 U/kg bleomycin. At 1-d post-BLM 1x10^6^ iCD103 cells generated from naïve CD11c-Cre+ AHR_Δex2_ mice were adoptively transferred via tail vein injection. Additionally, one group was administered 100 μg αCD4 neutralizing antibody at 1- and 7-d post bleomycin. Lungs were harvested 21-d post bleomycin and collagen content was quantified via hydroxyproline assay. **B through D)** iCD103 cells were generated as described. Cells were stimulated with 1 μM ODN2395 (TLR9 agonist), 10 μM RBN2397 (TiPARP inhibitor), or a vehicle control for 18 hours. Cells were harvested and RNA was analyzed for expression of IL-6 or TiPARP transcript via qRT-PCR. **E)** WT-B6 or AHR_Δex2_ mice were treated with 0.75 U/kg BLM, CH223191 (CH223) was administered from day 10- to 18-post BLM. Lungs were harvested at 21-day post BLM and expression of IL-6 transcript were quantified via qRT-PCR. **F and G)** Mice (n = 5 - 7 per group) were treated with 0.75 U/kg bleomycin or saline control (WT). Single cell suspensions were prepared from collagenase digested lung tissue at 7-days post bleomycin and CD11c+ cells were purified via magnetic bead isolation after which cells were pooled and treated overnight *ex-vivo* (n = 4 per group) with either 30μM CH223191 (CH223) or DMSO vehicle control. Expression of the indicated transcripts was analyzed via qRT-PCR. All experiments shown are representative of at least three independent experiments, statistical significance was determined via ANOVA (* = p < 0.05, ** = p < 0.01, *** = p < 0.001, **** = p < 0.0001).

The role of AHR in modulating cytokine responses is complex. Genes such as TiPARP induced through the canonical AHR pathway downregulate cytokine and interferon responses through post-translational modification of critical kinases such as TBK1 thus shutting down NFκB and IRF responses.(44–47) In contrast, AHR itself can directly bind NFκB and drive pro-inflammatory cytokine responses.(27, 48–50) Multiple TLRs, including TLR9, have been implicated in the progression of pulmonary fibrosis.(29, 51, 52) To address if DC IL-6 expression was dependent on cooperation between AHR and TLR signaling, we stimulated iCD103 DCs with the TLR9 agonist ODN2395. Cells generated from AHR_Δex2_ mice expressed over 2-fold more IL-6 in comparison to ODN2395-treated iCD103 cells from WT mice (Fig. 4B). Interestingly, while ODN treatment could stimulate expression of TiPARP in WT iCD103 DCs, this effect was abrogated in AHR_Δex2_ iCD103 DCs (Fig 4C). Together, these data indicate expression of TiPARP is dependent on canonical AHR signaling while expression of IL-6 is augmented by ncAHR signaling. We next treated iCD103 cells with TiPARP inhibitor RBN2397 and observed expression of IL-6 was increased by ∼2.5-fold when TiPARP was inhibited indicating production of TiPARP by canonical AHR signaling significantly inhibits non-canonical signaling and decreases production of inflammatory IL-6 (Fig. 4D). We next examined levels of IL-6 in BLM treated WT or AHR _Δex2_ mice that had been treated with the complete AHR inhibitor CH223191. In agreement with the increased IL-6 we observed in iCD103 DCs, AHR _Δex2_ mice exhibited increased expression of IL-6 both in whole lung and in CD11c+ cells isolated by magnetic bead purification from single cell lung digestions (Fig. 4E and 4F). Expression of IL-6 could be diminished by AHR inhibition (CCH223191) both at the whole lung and CD11c+ cell level indicating a direct effect of AHR in CD11c+ cells in potentiating IL-6 production in the fibrotic lung (Fig. 4E and 4F). Furthermore, we verified that expression of TiPARP was similarly down-regulated in lung CD11c+ cells from AHR _Δex2_ mice and thus conclude that the increased IL-6 observed is a product of the loss of anti-inflammatory canonical AHR signaling and a concomitant increase in pro-inflammatory non-canonical AHR signaling.

### DC ncAHR signaling potentiates IL-17 production from lymphocytes

Given that AHR_Δex2_ mice developed increased fibrosis and that ex-vivo generated DCs produced increased IL-6 in response to TLR9 stimulus, we next examined if AHR_Δex2_ mice would over-produce IL-17 in response to BLM. We generated lung single cell suspensions at 7-days post-BLM and analyzed expression of IL-17 in lymphocyte populations via flow cytometry. Flow cytometric analysis revealed AHR_Δex2_ mice displayed increases in multiple IL-17 producing lymphocyte populations including conventional Th17 cells, IL-17A producing gamma-delta T-cells, and putative IL-17A+ iNKT cells and ILC3s (Fig 5A quantified in 5B). Although we determined CD4+ lymphocytes were not responsible for increased fibrosis (Fig. 4A), we noted the largest increase in IL-17A expressing lymphocytes was in CD4- γδ TCR+ populations which would not be affected by CD4 depletion. AHR_Δex2_ mice expressed ∼5-fold more IL-17 at 21-days post-BLM (Fig. 5C).

**Figure 5.**
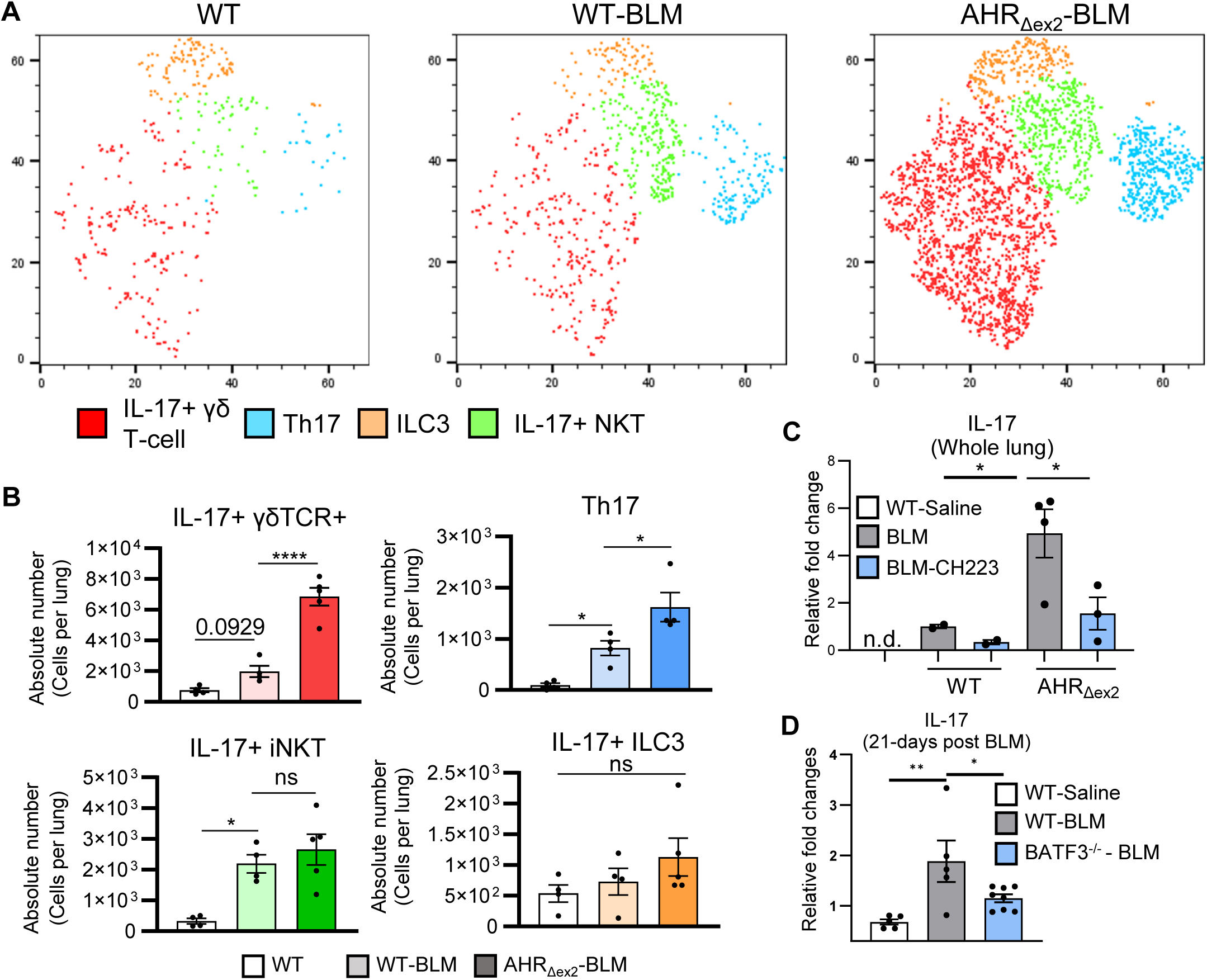
ncAHR signaling in CD11c+ cells drives IL-17 production in the lung. **A)** WT B6 or AHR_Δex2_ mice (n = 5 per group) were treated o.p. with 0.75 U/kg BLM. 7-days post-BLM lungs were harvested and lung leukocytes were analyzed by flow cytometry for the following populations Th17 (CD45+, Thy1.2+, CD3+, CD4+, IL-17A+), IL-17+ γδ t-cell (CD45+, Thy1.2+, CD3+, CD4-, γδ-TCR+, IL-17A+), IL-17+ ILC3 (CD45+, Thy1.2+, CD3-, CD4-, IL-17+), IL-17A+ iNKT (CD45+, Thy1.2+, CD4-, CD3+, γδ-TCR+, IL-17A+). **B)** Quantification of cell populations identified in A. **C)** WT-B6 or AHR_Δex2_ mice were treated with 0.75 U/kg BLM, CH223191 (CH223) was administered from day 10- to 18-post BLM. Lungs were harvested at 21-day post BLM and expression of IL-17 transcript was quantified via qRT-PCR. **D)** Mice (n = 5-8 per group) were treated with 0.75 U/kg BLM. 21-days post BLM treatment, lungs were harvested and expression of IL-17 transcript from whole lung was analyzed via qRT-PCR. All experiments shown are representative of at least three independent experiments, statistical significance was determined via ANOVA (* = p < 0.05, ** = p < 0.01, **** = p < 0.0001).

Production of both IL-6 and IL-17 was susceptible to complete inhibition of AHR via administration of CH223191 (Fig. 4E, 4F and Supplemental Fig. 2) indicating DCs and macrophages in CD11c-Cre / AHR_Δex2_ mice still had functional ncAHR signaling. We further verified the involvement of CD103+ DCs by examining expression of IL-17 in the lungs of BLM treated BATF3^-/-^ mice and found that loss of CD103+ DCs almost completely abrogated lung IL-17 responses (Fig. 5D).

### Fibroblasts from fibrotic lungs express kynurenine producing enzymes in response to TLR9 stimulus

Endogenous AHR ligands are largely produced as a result of the metabolism of tryptophan along the kynurenine pathway by enzymes such as IDO1 and TDO2. To determine what cells produce these enzymes in fibrotic lungs, we investigated data from two previously published single-cell RNAseq studies of IPF.(36, 37) Over 9000 stromal cells from 36 controls and 44 lungs were collectively analyzed and reclassified based on the markers and nomenclature described in Adams et al.(37) Though IDO1 was undetected, expression of TDO2 was observed upregulated specifically in IPF alveolar fibroblasts across both datasets (Fig. 6A and Supplemental Fig. 3). In addition to myeloid cells, TLR9 activation in fibroblasts has also been shown to be important in driving fibrosis.(29, 53) Thus, we next isolated fibroblasts from mouse lungs on d-21 post-Saline or BLM and subsequently treated these cells with ODN2395 to simulate an inflammatory / fibrotic environment *ex vivo*. Expression of both IDO1 and TDO2 increased in ODN2395-treated fibroblasts from BLM-treated mice but not in ODN2395-treated control fibroblasts (Fig. 6B and 6C). AHR was also upregulated in the same conditions (Fig 6D); however, TLR9 was upregulated with ODN2395 regardless of BLM treatment (Fig 6E). We further assayed for expression of the main enzymes involved in the production of kynurenine, i.e., IDO1 and TDO2, in whole lung following BLM treatment. Interestingly, expression of IDO1 was decreased at both 7- and 14-days post BLM (Fig. 7A); however, TDO2 expression was significantly increased at both time points (Fig. 7B) thus it appears that TDO2 is preferentially induced over IDO1 in pulmonary fibrosis *in vivo*. To address the importance of TDO2 during fibrosis we next treated groups of mice with the specific TDO2 inhibitor 680C91. Strikingly, inhibition of TDO2 significantly decreased both collagen deposition and expression of the myofibroblast marker alpha smooth muscle actin (αSMA) during BLM induced pulmonary fibrosis as well fibrotic pathology as assessed by trichrome staining of lung sections (Fig. 7C, 7D, and 7E).

**Figure 6.**
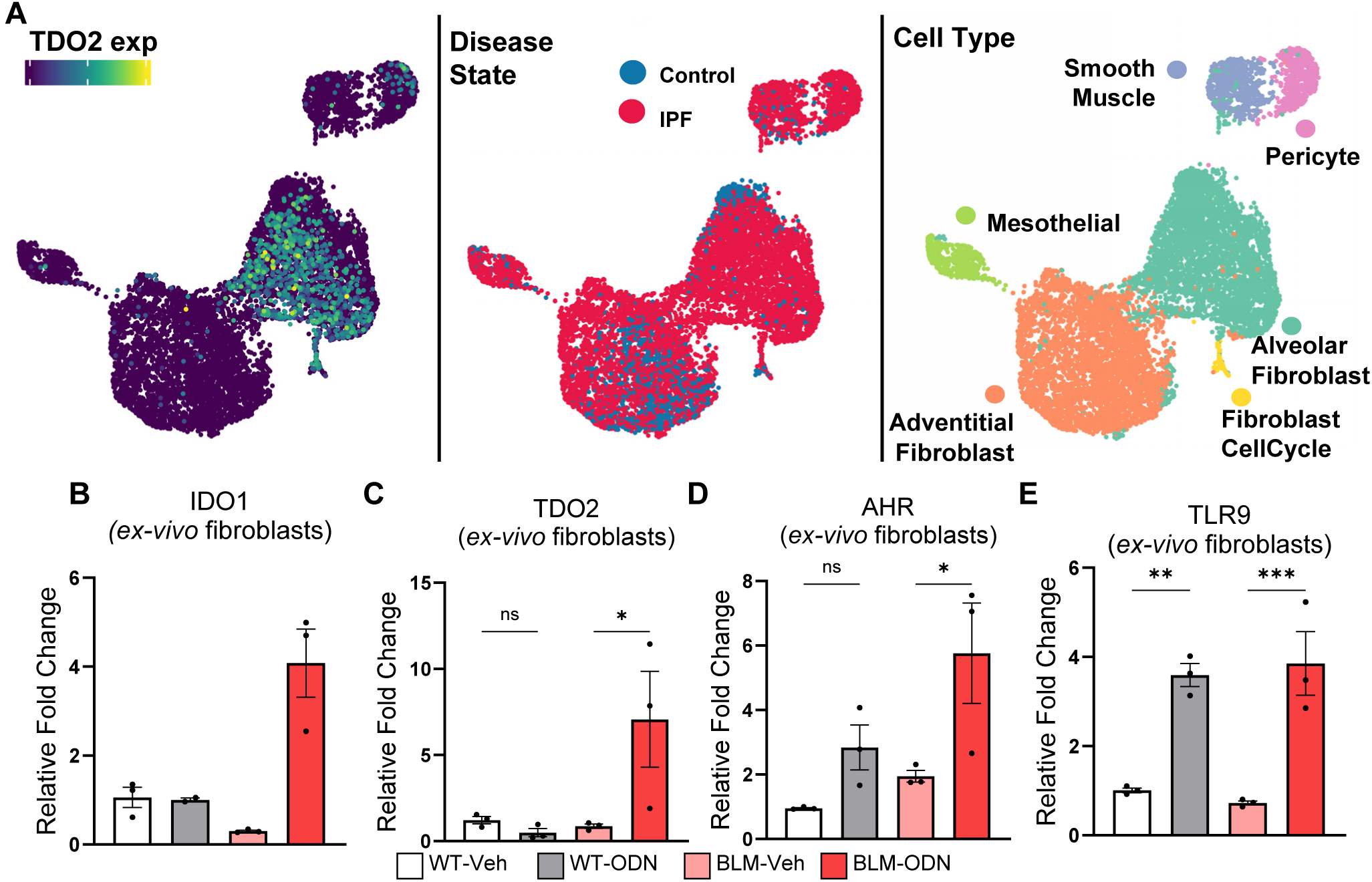
Fibroblasts isolated from fibrotic individuals express the kynurenine pathway enzymes IDO and TDO in response to TLR9 stimulation. **A)** Uniform manifold approximation and projections (UMAPs) of human lung stromal cells labeled by normalized expression of TDO2, cell type, and disease. **B through E**) Lung fibroblasts were cultured from groups of mice treated with BLM for 21-d or untreated as control. Fibroblasts were subsequently treated with the TLR9 agonist ODN2395 (ODN) or vehicle control (veh) for 18-h. Expression of the indicated transcript was analyzed via qRT-PCR. All experiments shown are representative of at least two independent experiments, statistical significance was determined via ANOVA (* = p < 0.05, ** = p < 0.01, *** = p < 0.001).

**Figure 7.**
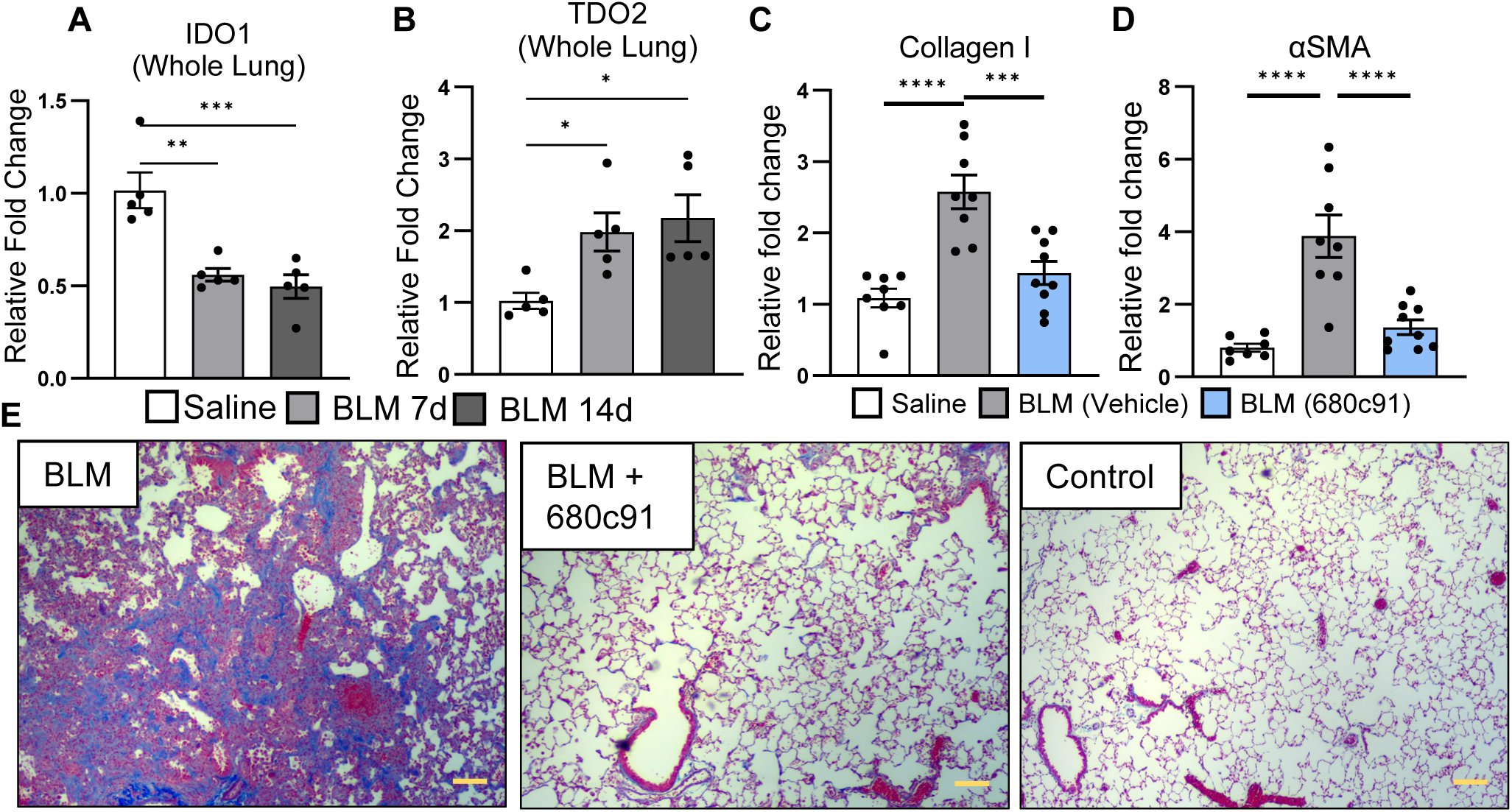
Inhibition of TDO2 rescues BLM induced fibrosis. **(A and B)** Groups of mice (n = 5 per group) were treated with 0.75 U/kg BLM or saline control for 7-14 days after which expression of the indicated transcript was analyzed in whole lung via qRT-PCR. **(C and D)** Group of mice (n = 8 – 9 per group) were treated with 0.75 U/kg BLM or saline control. Beginning on day 10 one group was treated with 15 mg/kg TDO2 inhibitor (680C91) daily by oral gavage until day 18 after which lungs were harvested and expression of the indicated transcript was assayed via qRT-PCR. **(E)** Representative histology (n =2 mice per group) showing assessment of collagen deposition by trichrome staining (blue staining) scale bar represents 100μm. All experiments shown are representative of at least two independent experiments, statistical significance was determined via ANOVA (* = p < 0.05, ** = p < 0.01, *** = p < 0.001).

## Discussion

### Role of DCs in regulating pulmonary fibrosis

The role of myeloid cells, especially DCs, in altering immune responses during development / exacerbation of pulmonary fibrosis is currently not well understood. Increased numbers of DC subsets infiltrate lungs of IPF patients and mice with pulmonary fibrosis.(8, 54–56) However, the study of DCs in pulmonary fibrosis has been overlooked due to publication of seminal work demonstrating that CD4+ and CD8+ lymphocytes, and by extension the DCs that prime those responses, are not important in mediating fibrotic effects.(57) Recently, it is appreciated that DCs can modulate immune responses outside of antigen presentation through production of pro-inflammatory / fibrotic cytokines. We, and others, have demonstrated that lung DCs can prime and activate pathogenic IL-17 responses through production of inflammatory cytokines such as IL-6 and TGF-β in innate lymphocytes as well as CD4+ T cells.(5, 58)

Here we demonstrate CD103+ DCs infiltrate the fibrotic lung and mediate production of pro-inflammatory / fibrotic cytokines through cooperation of AHR and pro-inflammatory TLR signaling. We noted significant accumulation of CD103+ DCs in lungs of BLM-treated mice beginning in the early inflammatory phase (7-days post-treatment) and continuing into the fibrotic phase (21-days post treatment, Fig. 1F). Loss of CD103+ DCs in BATF3^-/-^ mice was protective indicating a pathogenic role for lung resident DCs (Fig. 1B – 1E). In agreement with our previously published study(4), CD103 cells that infiltrate the fibrotic lung express high levels of AHR (Fig. 1F). Canonical AHR signaling is immuno-suppressive and AHR target genes such as TiPARP can constrain production of interferons and pro-inflammatory cytokines via ADP-ribosylation of activators of NF-κB signaling such as TBK1.(44–47) Alternatively, ncAHR signaling is well characterized to bind NF-κB subunits like RelA and RelB and enhance production of pro-inflammatory cytokines.(27, 48–50) We noted increased expression of IL-6 following BLM both in the whole lung and from isolated CD11c+ cells which correlated with increased expression of AHR (Fig. 1); however, expression of the AHR inducible gene product, Cyp1b1 was significantly decreased (Fig. 1H). Thus, even though CD11c+ cells express AHR and there is abundant AHR ligand produced in fibrotic lungs (Fig. 2A), these cells do not activate canonical AHR signaling and instead activate pro-inflammatory ncAHR signaling in the BLM treated lungs.

### Role of ncAHR / NF-Kb and TIPARP in modulating cellular cytokine responses

The role of tryptophan metabolism and AHR signaling in mediating pulmonary fibrosis is unclear. Kynurenine / tryptophan ratios increase in serum and lungs of patients with interstitial pneumonia and treatment with anti-fibrotic Pirfenidone significantly decreased levels of kynurenine suggesting a positive correlation between tryptophan metabolism, AHR activation, and development of fibrosis.(12, 59) More recently, activation of AHR through exogenous administration of AHR ligand FICZ ameliorated pulmonary fibrosis presumably by augmenting numbers of FoxP3+ regulatory T-cells, and decreasing IFNγ+ CD4+ T-cells and IL-17A+ γδ+ T-cells.(60) However, this study did not look specifically at AHR signaling in pulmonary immune cells nor did it demonstrate if FICZ is produced during fibrosis. Furthermore, mice were treated directly after administration of BLM on days 0, 1, and 2 making it difficult to determine direct effects of pulmonary AHR signaling in inhibiting fibrosis versus modulating early lung injury post-BLM. We detected significantly elevated kynurenine from 7 – 14 days post-BLM corresponding to the early fibrotic phase of BLM induced fibrosis (Fig. 2A). In contrast to previous work, complete inhibition of AHR signaling via administration of CH223191 reduced fibrosis when started late in the progression of disease (10-days post-BLM) indicating a role for pulmonary AHR signaling in promoting fibrogenesis (Fig. 2B – 2C).

We generated mice that lacked canonical AHR signaling in CD11c+ cells including pulmonary DCs and macrophage (Fig. 3A and Supplemental Fig. 1). Interestingly, while DCs produced from these mice lacked canonical AHR signaling (Fig. 3B), we still detected AHR at the transcript and protein level (Fig. 3C – 4E). Furthermore, sequencing cDNA generated from AHR_Δex2_ DCs indicated an alternative start codon is formed when exon 2 is deleted (Supplemental Fig. 1A). Together, these data indicated that a truncated AHR product still mediated ncAHR interactions. When we administered BLM to AHR_Δex2_ mice we noted more fibrosis that was still AHR-dependent (Fig. 3F and 3G) indicating loss of canonical AHR signaling in CD11c+ cells resulted in increased ncAHR signaling that exacerbated pulmonary fibrosis.

To determine if increased fibrosis seen in AHR_Δex2_ was intrinsic to DCs we adoptively transferred ex-vivo generated iCD103 DCs from AHR_Δex2_ mice into BLM treated B6 mice and noted more fibrosis when DCs from CD11c-Cre+ / AHR_Δex2_ were transferred in comparison to either Cre-cells (data not shown) or non-transferred mice (Fig. 4A). We further show AHR can synergize with TLR9 activation and promote expression of IL-6 in DCs; however, expression of canonical AHR genes that limit cytokine production, e.g., TiPARP is lost in AHR_Δex2_ DCs (Fig 4B – 4D). Thus, BLM-induced fibrosis results in increased expression of AHR; however, the signaling is shunted towards non-canonical signaling resulting in increased expression of pro-inflammatory cytokines. It is currently unclear why ncAHR signaling is promoted over canonical signaling in fibrotic lungs, however, several transcription factors such as HIF1α which are up-regulated during fibrogenesis bind to the canonical AHR partner ARNT (HIF1β).(61) Thus, it is plausible that dimerization of HIF1α-ARNT may reduce the available pool of ARNT which promotes AHR-NFκB dimerization shifting the balance of AHR signaling from canonical, anti-inflammatory, to non-canonical, inflammatory, signaling which drives excessive fibrogenesis.

### Role of IL-6 in promoting profibrotic IL-17 responses

The role of the IL-6 – IL-17 axis is appreciated in the progression of pulmonary fibrosis. Depletion of either IL-6 or IL-17 is protective for BLM-induced fibrosis and we have previously published that IL-17 can act directly on fibroblasts to promote collagen production and fibroproliferation.(5, 62–64) Even in IPF patients, expression of IL-6 and IL-17 isoforms are associated with more rapid time to death or lung transplantation (Supplemental Fig. 3).

In agreement with this we isolated fibroblasts from control or BLM-treated mice and treated with the TLR9 agonist, ODN2395. TLR9 activation upregulated expression of both IDO1 and TDO2 in BLM fibroblasts but not control (WT) fibroblasts, we corroborated expression of TDO2 in IPF patient myofibroblasts, however, we did not detect appreciable expression of IDO1 in human myofibroblasts and overall expression of IDO1 was shown to decrease in the lungs of fibrotic mice while expression of TDO2 increased (Fig. 6A – 6C, 7A and7B). Treatment of mice with the TDO2 inhibitor 680C91 ameliorated fibrosis, reducing collagen production, and decreasing expression of the myofibroblast marker αSMA (Fig. 7C, 7D, and 7E). Furthermore, we noted increased expression of AHR and TLR9 itself following TLR9 stimulation in BLM fibroblasts indicating that AHR plays a fundamentally pro-fibrotic role (Fig. 6D – 6E). However, In direct contrast to this, a previous study found that treatment of fibroblasts with the AHR activating compound ITE inhibited TGFβ induced myofibroblast differentiation and collagen production.(28) The exact cause for the discrepancy between these two studies will require further investigation, however, it is plausible that induction of AHR signaling in the context of inflammatory stimulus, e.g., TLR stimulation as would happen in the fibrotic lung behaves fundamentally different than in an isolated *ex vivo* condition. Expression of TLR9 is associated with rapid progression of IPF in patients and loss of TLR9 signaling is protective from BLM-induced fibrosis.(29, 51, 52) Furthermore, fibroblasts isolated from IPF patients that had more robust activation profiles when exposed to TLR9 agonist correlated with increased AHR activation in matched myeloid cells.(30) Together, these studies indicate that a fibrogenic circuit is established between TLR9+ fibroblasts and AHR+ myeloid cells that is important in progression of pulmonary fibrosis through modulation of pro-fibrotic cytokines like IL-6 and IL-17 (Working model in Fig. 8). More work is needed to fully understand the switch to ncAHR signaling during fibrogenesis and to understand the spatial crosstalk between fibroblasts and DCs. However, given the need for new treatments for IPF patients, drugs that modulate kynurenine pathway enzymes, i.e., IDO1 and TDO2, or AHR directly represent an exciting possibility with untapped therapeutic potential.

**Figure 8.**
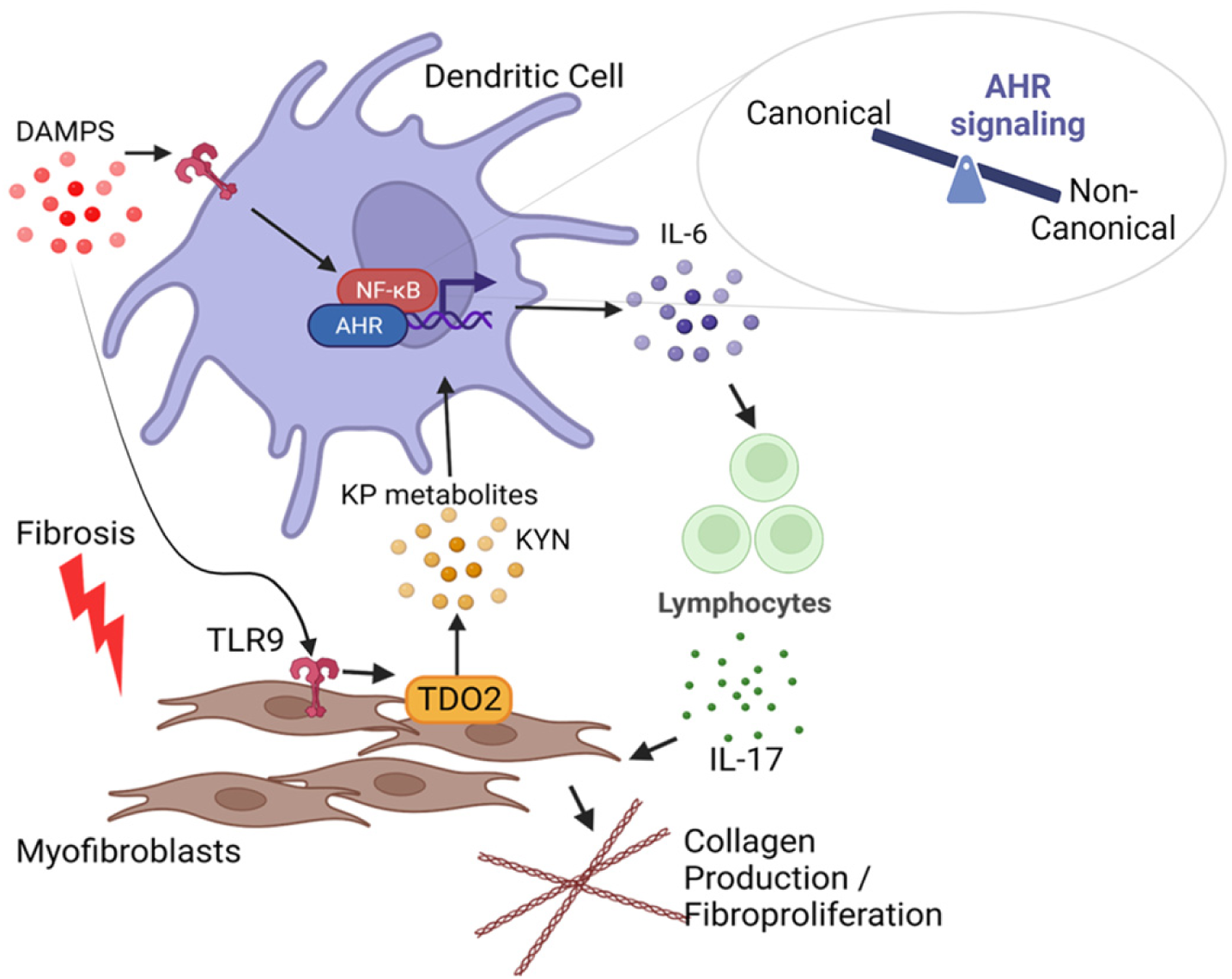
Working model for how fibroblast DC crosstalk contributes to fibrosis. Myofibroblasts express the kynurenine pathway enzyme TDO2 in response to TLR9 stimulation. Kynurenine pathway metabolites active ncAHR signaling in DCs which, in cooperation with inflammatory TLR singling, augments production of IL-6 resulting in production of IL-17 from lymphocytes. Fibroblasts can be directly activated by IL-17 resulting in collagen production and fibroproliferation.

