## Supplemental Figures for "Dendritic Cell – Fibroblast Crosstalk via TLR9 and AHR Signaling Drives Lung Fibrogenesis"

### Supplemental Figure 1

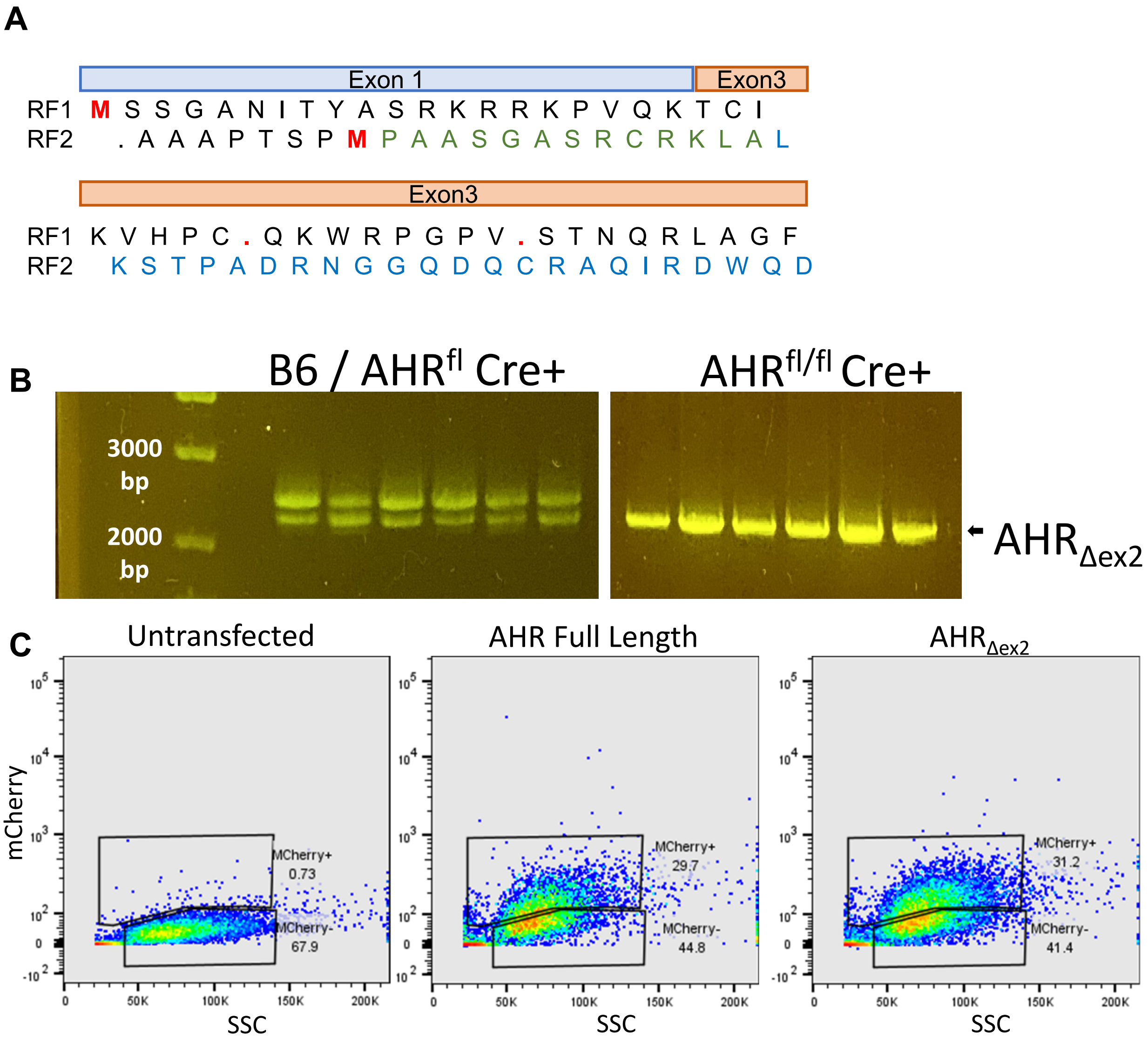


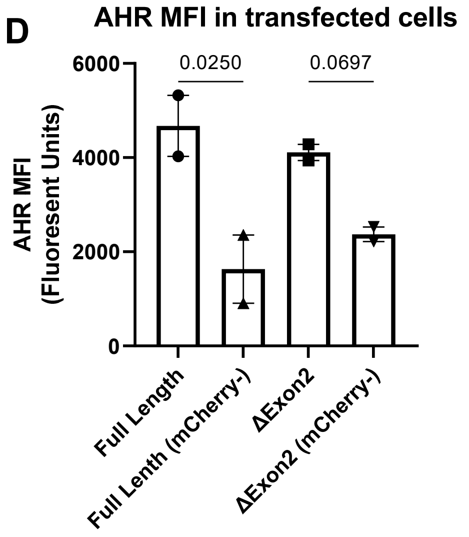


**Supplemental Figure 1. Generation of CD11c-Cre / AHR$^{fl/fl}$ mouse (AHR$_{\Delta ex2}$ ) and characterization of a truncated AHR product lacking exon 2.** (A)Schematic detailing the exon1-3 fusion that is produced in AHR$_{\Delta ex2}$ mice. Deletion of AHR exon 2 shifts the open reading frame from RF1 to RF2 (RF1 = original ORF, RF2 = alternative ORF) revealing an alternative start codon which results in translation of a 14 amino acid leader sequence (green text) and exons 3 – 10 in frame (blue text). (B) PCR of AHR CDNA from either B6 / AHR$^{fl}$ or AHR$_{\Delta ex2}$. (C and D) NIH 3T3 cells were transfected with the indicated clone and analyzed for mCherry and AHR co-expression by flow cytometry in panel C or expression of AHR in panel C.

### Supplemental Figure 2

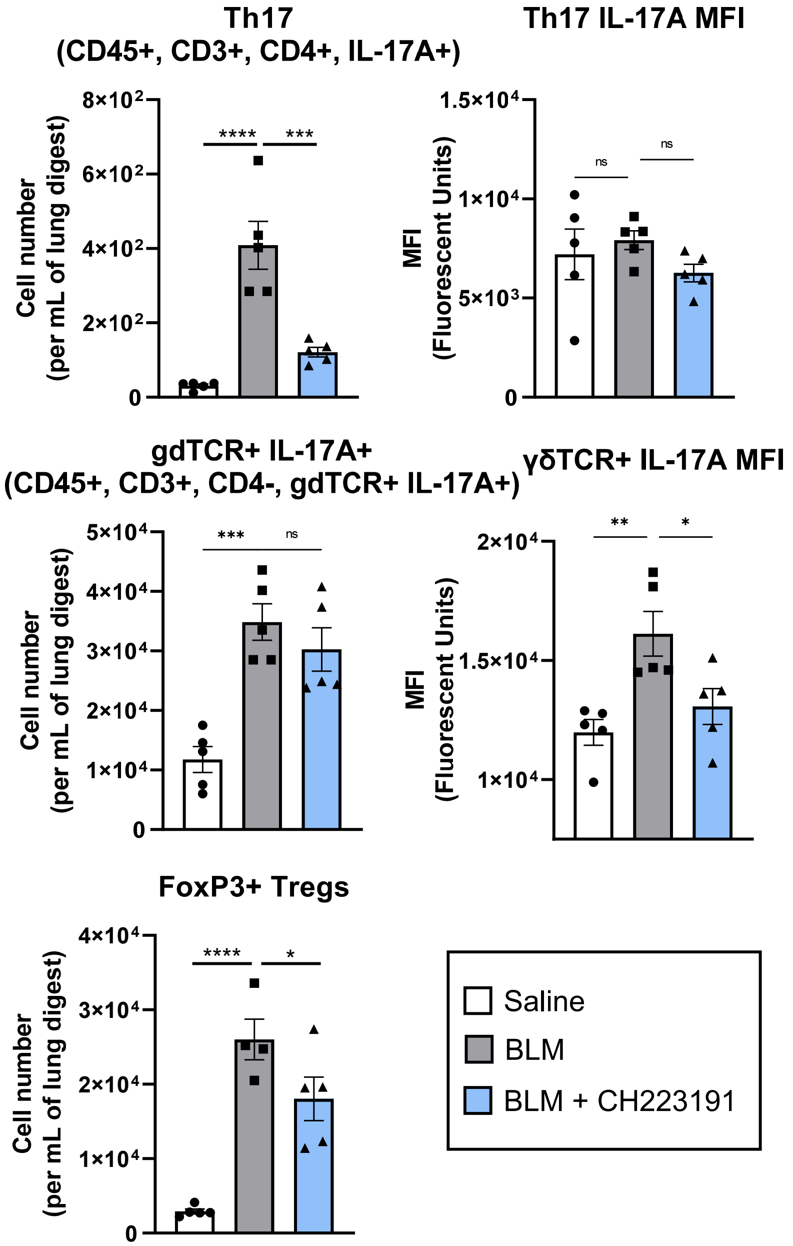


**Supplemental Figure 2. AHR inhibition reduces IL-17 production in Th17 and γδ T-cells.** Mice (n  -5 mice per group) were treated with BLM and lungs were harvested at 7 dpi. Flow cytometry was conducted on single cell suspensions to identify the following lymphocyte populations Th17 (CD45+, CD3+, CD4+, IL-17A+), IL-17 expressing γδ T-cells (CD45+, CD3+, CD4-, γδ TCR+, IL-17A+), and FoxP3+ Tregs (CD45+, CD3+, CD4+, FoxP3+).

### Supplemental Figure 3

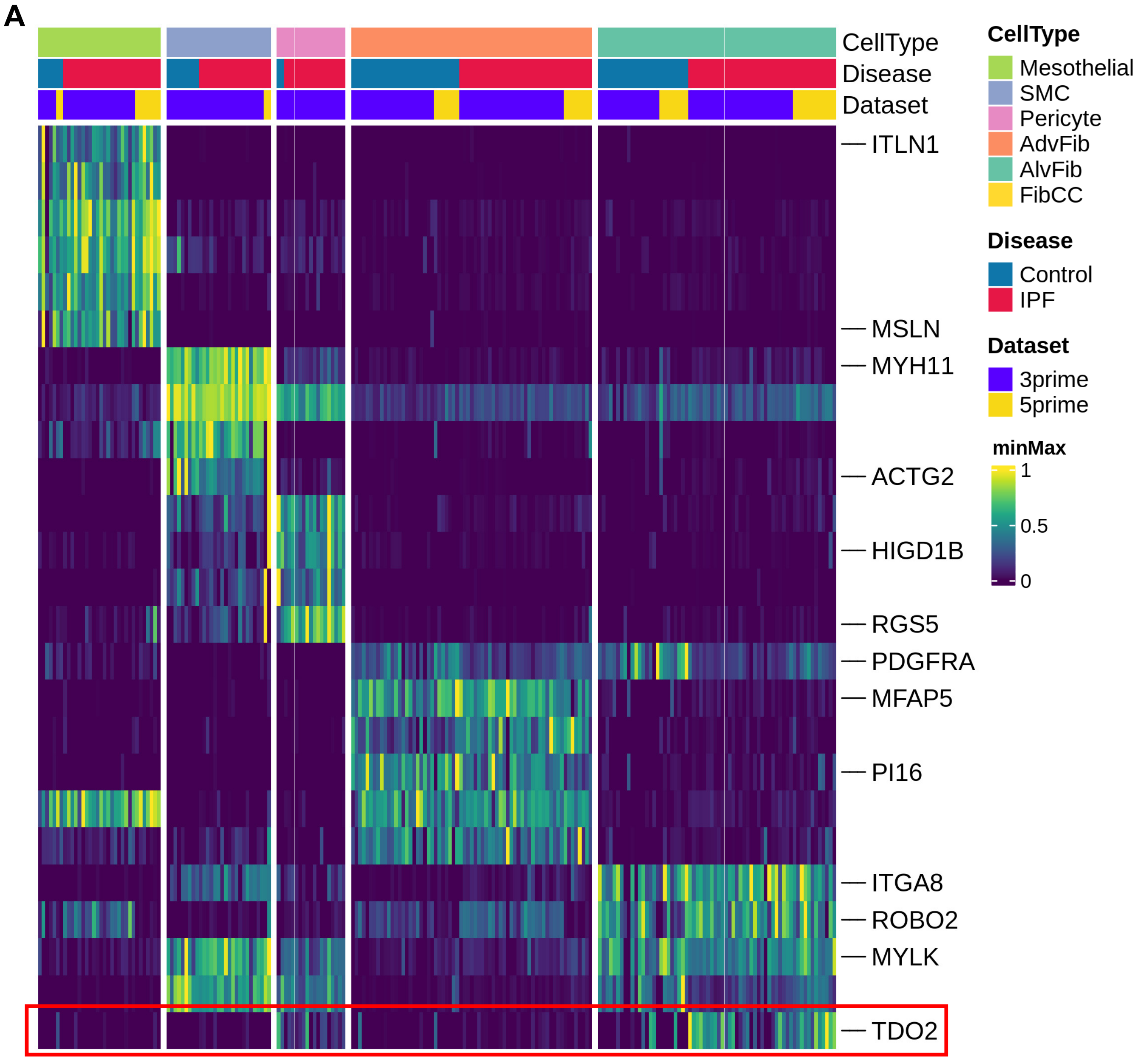


**Supplemental Figure 3. Myofibroblasts express TDO2 in the fibrotic lung and expression is consistent across multiple datasets. A)** Heatmap of gene expression values across human lung stromal cells. Each column represents the average normalized expression of one cell type and one subject. Within each dataset, expression values for each gene are min-max scaled to the range of 0-1.

### Supplemental Figure 4

**A**

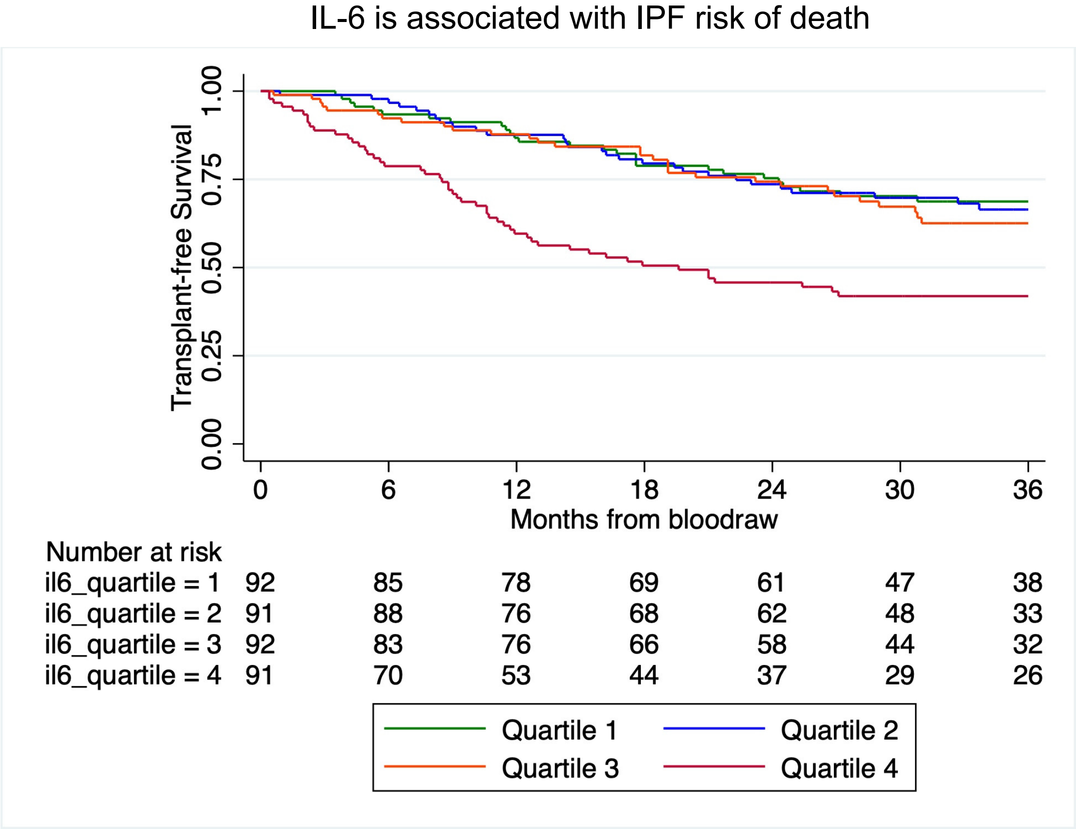


**B**

| Risk of death or lung transplant within 36-months of blood draw | | | |
|---|---|---|---|
| **Biomarker** | **HR** | **95% CI** | **p-value** |
| IL6 | 1.41 | 1.24-1.61 | <0.001 |
| IL17a | 0.98 | 0.87-1.12 | 0.83 |
| IL17c | 1.26 | 1.08-1.48 | 0.004 |
| IL17d | 0.55 | 0.36-0.83 | 0.005 |
| IL17f | 1.26 | 1.06-1.50 | 0.007 |

**Supplemental Figure 4. Plasma concentrations of IL-6 and IL-17 isoforms are associated with risk of death or transplant in IPF patients. A)** Survival curve showing correlation between IL-6 expression and transplant free survival in a cohort of IPF patients (n = 366). **B)** Table showing hazard ratios for IL-6 and Il-17 isoforms in IPF patients.

### Supplemental Table 1

| subset | CD45 | CD11b | CD11c | CD24 | CD64 | MHCII | SiglecF | CD103 | Ly6g | Ly6c | Thy1.2 |
|---|---|---|---|---|---|---|---|---|---|---|---|
| cDC1 | + | - | + | + | - | + | - | + | - | - | - |
| cDC2 | + | + | + | + | - | + | - | - | - | - | - |
| Alv. Macs | + | - | + | - | + | + | + | - | - | - | - |
| Int. Macs | + | + | + | - | + | + | - | - | - | - | - |
| T.R. Monos | + | + | + | - | - | - | - | - | - | - | - |
| Inf. Monos | + | + | - | - | - | - | - | - | - | + | - |
| Lymphocytes | + | - | - | - | - | - | - | - | - | - | + |
| B-Cells | + | - | - | + | - | + | - | - | - | - | - |
| Granulocytes | + | + | - | + | - | - | - | - | + | - | - |

**Supplemental table 1. Markers used for identification of lung leukocyte populations by flow cytometry.**
